# Enhanced epithelial to mesenchymal transition and chemoresistance in advanced Retinoblastoma tumors is driven by miR-181a

**DOI:** 10.1101/2022.07.25.501381

**Authors:** Vishnu Suresh Babu, Anadi Bisht, Ashwin Mallipatna, Deepak SA, Gagan Dudeja, Ramaraj Kannan, Rohit Shetty, Stephane Heymans, Nilanjan Guha, Arkasubhra Ghosh

## Abstract

Advanced retinoblastoma (Rb) tumors can infiltrate distant tissues and cause a potent threat to vision and life. Through transcriptomic profiling, we discovered key epithelial to mesenchymal transition (EMT) and chemotherapy resistance genes at higher expression levels in advanced Rb tumors. Rb-/- tumor cells acquire metastasis-like phenotype through the EMT program that critically contributes to chemoresistance. We demonstrate that prolonged chemo-drug exposure in Rb cells elicits an EMT program through *ZEB1* and *SNAI2* that further acquires therapeutic resistance through cathepsin L and *MDR1* mediated drug efflux mechanisms. Further, 16 significantly differentially expressed miRNAs were identified in patient tumors, of which miR-181a-5p was significantly reduced in advanced Rb tumors and associated with altered EMT and drug resistance genes. Enhancing miR-181a-5p levels in Rb-/- cells and Rb-/- chemo-resistant sublines controls EMT transcription factors *ZEB1* and *SNAI2* and halts the transition switch, thereby reversing drug resistance. We thus identify miR-181a-5p as a potential therapeutic target for EMT triggered drug-resistant cancers that can halt their invasion and sensitize them to low dose chemotherapy drugs.

**Graphical Abstract:** 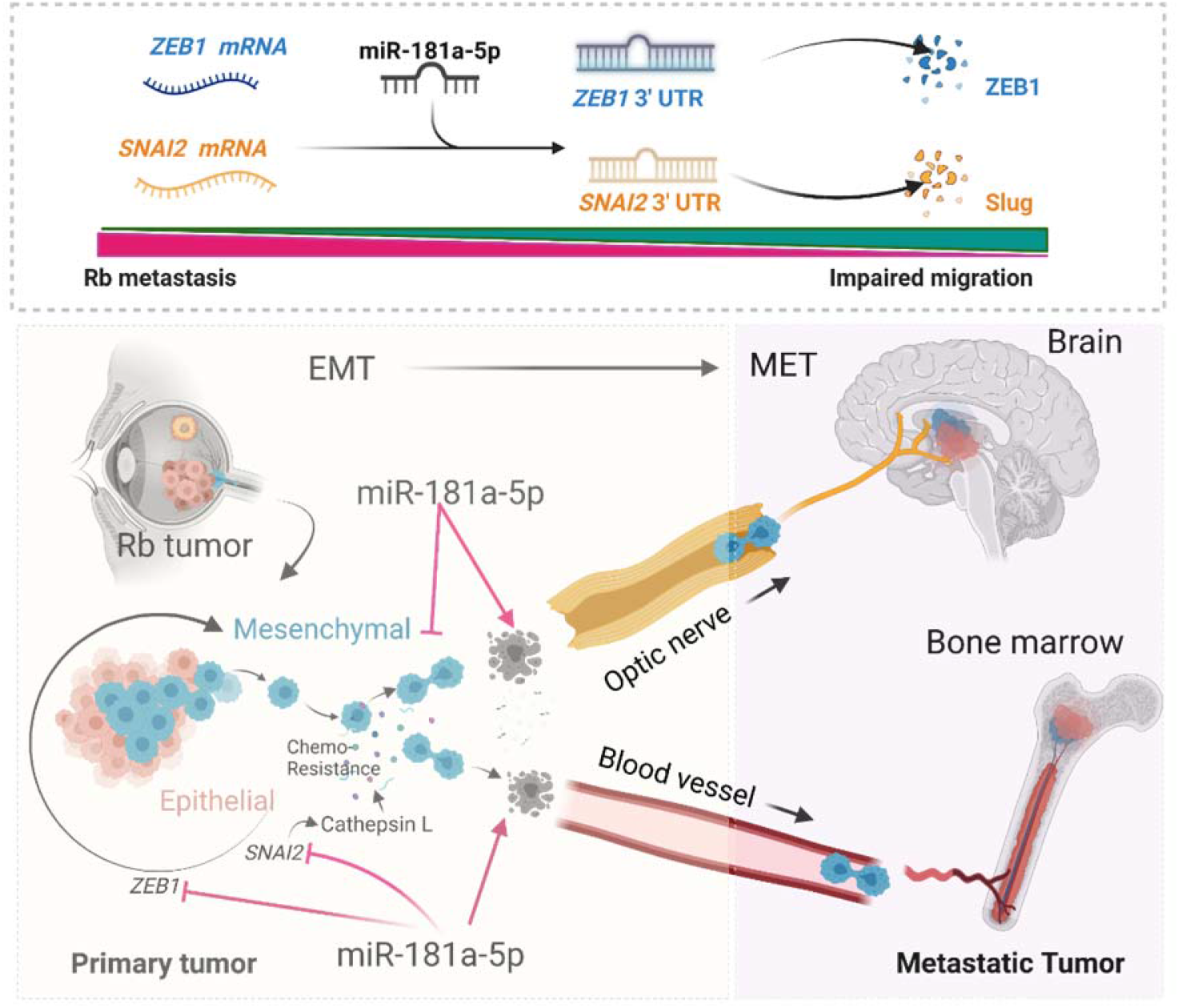

## Introduction

Retinoblastoma (Rb) is the most common intraocular malignant tumor in children. Managing intraocular Rb tumors by efficient diagnosis, genetic screening and clinical procedures (Ancona-Lezama et al., 2020; Gupta et al., 2021) help achieve excellent survival rates worldwide. However, metastatic retinoblastoma is still a major concern in many countries(Honavar et al., 2017; Vempuluru et al., 2021) (Canturk et al., 2010). Rb tumors that grow rapidly have sufficient feeder arteries and drainage veins, characterized by the presence of multifocal yellowish white tumor mass with floating subretinal or vitreous cancer seeds(Shields & Shields, 2010). If neglected or untreated, advanced Rb tumors demonstrate massive choroidal invasion (Bosaleh et al., 2012) and metastatic spread, primarily through optic nerve (Shields et al., 1994) and sclera (Rootman et al., 1978), to regional lymph nodes, central nervous system (CNS) and bone marrow(Finger et al., 2002) causing potent threat not only to vision, but to life of the child as well. Clinically, to manage metastatic Rb tumors, an intensive multimodal approach incorporating high dose systemic, intra-arterial, peri-orbital chemotherapy regimens involving carboplatin, etoposide and cyclophosphamide followed by radiation are used currently (Namouni et al., 1997). However, advanced tumors often evolve during successive chemotherapy cycles and develop resistance to anticancer therapeutics, diminishing the efforts of the clinical management procedures (Chan et al., 1991; Shields et al., 2003). Advanced Rb tumors upon prolonged chemo-drug exposure, increases the expression of ATP binding cassette (ABC) transporter pathway genes like *MDR1* and *MRP1* to confer resistance by chemo-drug efflux mechanism (Wilson et al., 2006). Metastatic tumors acquire chemotherapy resistance through trans-differentiation initiated by the epithelial to mesenchymal transition (EMT) program in different cancers (Choi et al., 2019) (Saxena et al., 2011). EMT program begins with the loss of epithelial phenotypes by downregulation of E-cadherin and tight junction adhesion molecules. The differentiated cancer cells transit to mesenchymal phenotype with an invasive dedifferentiated characteristic, which can coincide with acquiring chemo-drug resistance properties.

MicroRNAs (miRNA) are small non-coding single strand RNAs that have emerged as important modifier of plethora of biological pathways including cancers(Lin & Gregory, 2015). They modify gene expression by using the RNA-induced silencing complex (RISC) that bind to the 3’ untranslated region (UTR) or less frequently 5’ UTR region of the mRNA and cause translational repression. Emerging evidences points out the role of miRNAs in controlling EMT transcription factors and signaling pathways to regulate metastatic dissemination in different cancers (Diaz-Lopez et al., 2014). In Rb tumors, increased expression of miR 17-92 cluster (Kandalam et al., 2012), miR-25-3p (Wan et al., 2020) & miR200c (Shao et al., 2017) were found to regulate EMT mediated high invasion and migration of Rb cells in-vitro, thus supporting the role of EMT in Rb metastasis. However, the mechanistic link between miRNA, EMT and drug resistance in Rb patient tumors remain obscure.

In the present study, we profiled miRNA and mRNA signatures simultaneously in the same set of advanced and non-advanced Rb tumors compared to age matched healthy pediatric retina. Such a coordinated analysis of expression networks in the same set of tissues and controls enabled the discovery of co-regulated miRNA and mRNA targets relevant to Rb stage. Among the many dysregulated genes and miRNAs, we chose to validate and investigate the functional role of miR-181a-5p on the enhanced EMT and drug resistance pathways in advanced Rb subjects.

## Results

### Transcriptomic profiling identifies differentially regulated miRNA’s, EMT, and drug-resistant genes in Rb tumor subtypes

To investigate alterations to miRNA mediated regulation in Rb tumors, we generated the miRNA profile from patient tumors. The Rb tumor sample cohort (Table 1) comprised of enucleated tumor tissues of five advanced (defined by AJCC staging-cT3(Mallipatna et al., 2017), IIRC-Group E (Linn Murphree, 2005) and four non advanced (defined by AJCC staging-cT2, IIRC-Group D) subjects. Two age matched pediatric retina (age range from 2-3 months) obtained from donors without ocular complications were used as controls. We identified sixteen highly significant, differentially regulated miRNAs unique to Rb tumors (*P*<0.05, FC>2) compared to pediatric retina (Figure 1A). Notably, miR-181a-5p and miR-3653 were significantly downregulated in advanced Rb compared to non-advanced Rb tumors (Figure 1B). We applied KEGG pathway enrichment analysis to the miRNAs data obtained by microarray and identified miRNAs regulating genes belonging to cell cycle pathway, EMT program, drug resistance and pathways in cancer (Figure 1C). We performed RT-PCR validation experiments in a secondary cohort (Table S1) comprising of eight Rb tumor tissues (4 advanced, 4 non advanced) and four pediatric retina controls, that confirmed the downregulated of miR-181a-5p in Rb tumors, with significant downregulation in advanced subjects (*P*<0.001) (Figure 1D). RT-PCR quantification of miR-331-3p, miR-574-5p and miR-1290 in advanced and non-advanced Rb tumors corroborated with the expression profiles identified in miRNA microarray (Figure S1A, B, C). The findings prompted us to elucidate the EMT and drug resistance signatures in advanced and non-advanced Rb tumors. We have previously performed total mRNA profiling using gene expression microarray in the primary Rb cohort (Babu et al., 2022). We identified distinct clustering of differentially regulated EMT and drug resistance genes in advanced (Figure 1E) and non-advanced Rb tumors (Figure 1F) compared to pediatric controls (*P*<0.05, FC>2). Notably, EMT transcription factors like *ZEB1* (FC=92, *P*<0.05), *SNAI2* (FC=5.57, *P*<0.05) and drug resistance genes like *ABCB1* (MDR1) (FC=5.84, P<0.05), *CTSL* (Cathepsin L) (FC=20.03, *P*<0.05) were significantly upregulated in advanced tumors (Figure 1G). However, ZEB1 (FC=77.2, *P*<0.05), SNAI2 (FC=3.32, *P*<0.05), *ABCB1* (FC=4.4, *P*<0.05) were low and *CTSL* (FC= −3.8, *P*<0.05) expressions was significantly downregulated in non-advanced Rb tumors. RT-PCR validations of these genes in a secondary cohort confirmed the findings of the microarray (Figure 1H-K).

**Figure 1:**
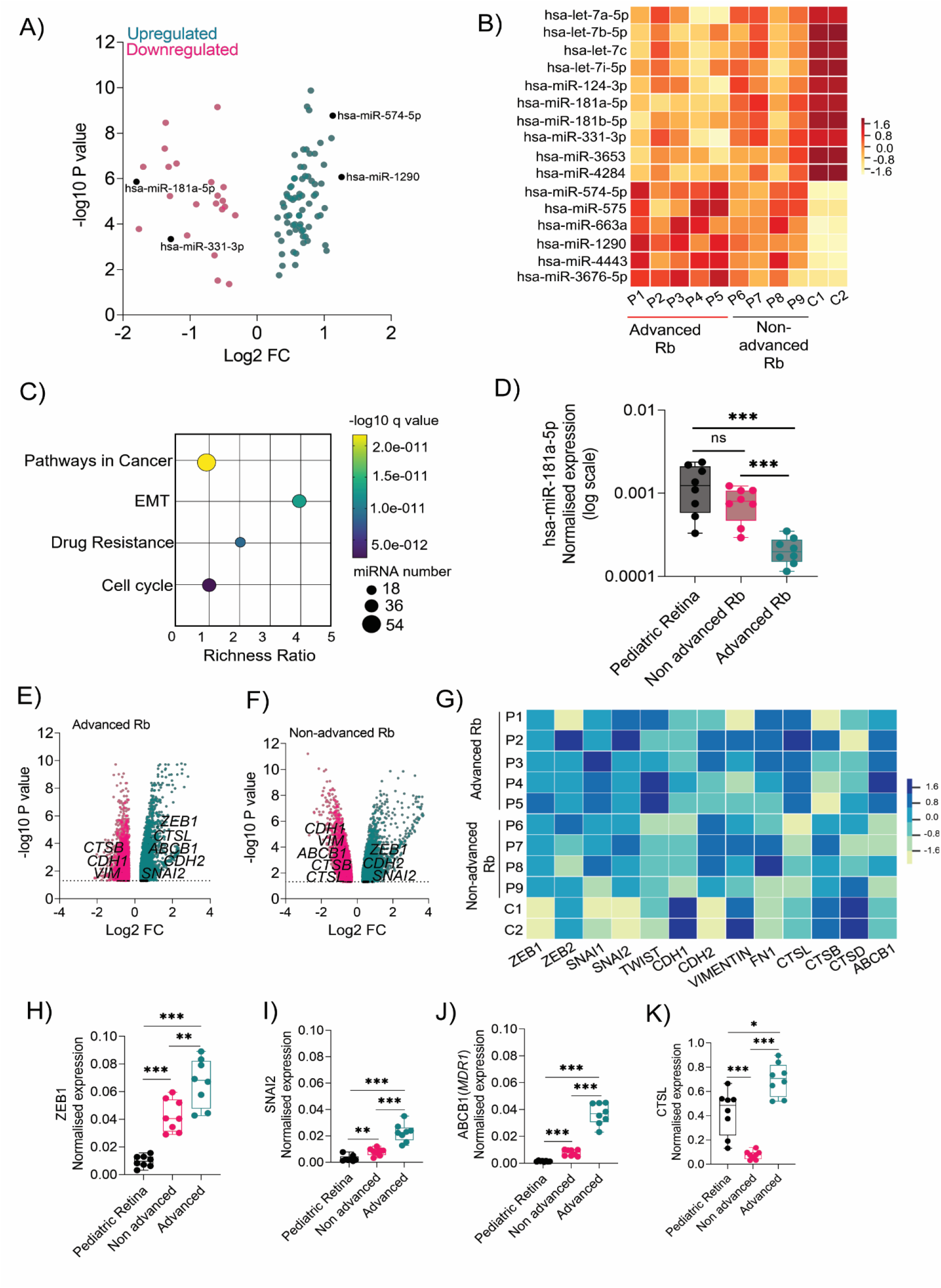
Transcriptomic profiling identifies differentially regulated miRNA’s, EMT, and drug-resistant genes in Rb tumor subtypes. (A) Volcano plot showing differentially regulated miRNA’s in Rb subjects (n=9) compared to pediatric retina (n=2) identified using microarray. (B) Heatmap showing differential expression of miRNAs in 9 Rb subjects and 2 pediatric controls identified using microarray. (C) Bubble scatter plot showing top enriched KEGG pathways regulated by miRNAs in Rb tumors. (D) RT-PCR results showing normalised expression of miR-181a-5p in control retina (n=4), advanced Rb (n=4) and non-advanced Rb (n=4). Volcano plot showing differentially regulated EMT and chemotherapy resistant genes identified using microarray in (E) Advanced Rb tumors (F) Non-advanced Rb tumors. (G) Heatmap showing expression of EMT and chemotherapy resistant genes in 9 Rb subjects and 2 pediatric controls. RT-PCR showing normalized expression of (H) *ZEB1* (I) *SNAI2* (J) *ABCB1* and (K) *CTSL* in control pediatric retina (n=4), advanced (n=4) and non-advanced (n=4) Rb tumors. Values represents mean ± s.d. Two tailed Mann-Whitney was used for statistical analysis. *p < 0.05, **p < 0.01, ***p < 0.001, ****p < 0.0001.

**Table 1:** Clinical and histopathological details of samples used in the study.

| ID | Sex | Laterality | Age at presentation | Clinical Risk | ICRB Group | AJCC Staging |
| --- | --- | --- | --- | --- | --- | --- |
| P1 | M | Bilateral | 15 months | Advanced | Group E | cT3b |
| P2 | F | Unilateral | 20 months | Advanced | Group E | cT3b |
| P3 | M | Unilateral | 24 months | Advanced | Group E | cT3a |
| P4 | F | Bilateral | 4 months | Advanced | Group E | cT3b |
| P5 | M | Bilateral | 30 months | Advanced | Group E | cT3b |
| P6 | F | Bilateral | 21 months | Non-advanced | Group D | cT2b |
| P7 | F | Unilateral | 28 months | Non-advanced | Group D | cT2b |
| P8 | M | Unilateral | 20 months | Non-advanced | Group D | cT2b |
| P9 | M | Unilateral | 21 months | Non-advanced | Group D | cT2a |
| Control 1 | F | NA | 3 months | Cardiac Arrest (no ocular complications) | NA | NA |
| Control 2 | F | NA | 2 months | Multiple organ dysfunction ( No ocular complications) | NA | NA |

### Validation of epithelial to mesenchymal transition (EMT) and chemo-drug resistance proteins in Rb tumors and their interaction with miR-181a-5p

Immunofluorescence analysis on FFPE specimens of Rb tumors detected strong ZEB1 and Cathepsin L positivity in advanced Rb tumor tissues compared to non-advanced Rb (Figure 2A, B & C). However, we observed cathepsin L positivity in photoreceptor layers of control tissues, in line with its known lysosomal functions in the retina (Sharif et al., 2019) unlike its metastatic role in cancers (Dykes et al., 2019). Notably, advanced Rb tumors demonstrated high expression of N-cadherin and low expression of E-cadherin (Figure 2D, E & F) indicative of the EMT process (Loh et al., 2019). RT-PCR validations further confirmed the downregulation of *CDH1* (E-cadherin) in advanced and non-advanced tumors (*P*<0.001) (Figure S1D) while *CDH2* (N-cadherin) maintained an elevated expression profile in advanced tumors (*P*<0.001) (Figure S1E). We also detected increased MDR1 levels in advanced tumors compared to the non-advanced tumors and controls, indicating resistance to drug therapy (Figure S1F). For comprehensive functional analysis of miRNAs, we developed a miRNA-target interaction network map using miRNet by integrating the microarray data with microRNA databases like miRanda, miRbase and TargetScan. Out of the sixteen miRNAs identified in Rb tumor microarray, nine miRNAs were associated with different cancer related pathways while five miRNAs were predicted to specifically regulate EMT pathway genes in the interaction map. We identified miR-181a-5p as a regulator of EMT transcription factors like *ZEB1* and *SNAI2*, while miR-124-3p was identified as a regulator of EMT facilitators like *CDH1* (E-cadherin) and *CHD2* (N-cadherin) (Figure 2G). However, miR-3653 did not display any interaction with KEGG identified enriched pathways (Figure 2G). We speculate that advanced tumors maintain high expression of EMT and drug resistance genes due to low expression of miR-181a-5p, thus promoting invasion and metastasis (Figure 2H, I).

**Figure 2:**
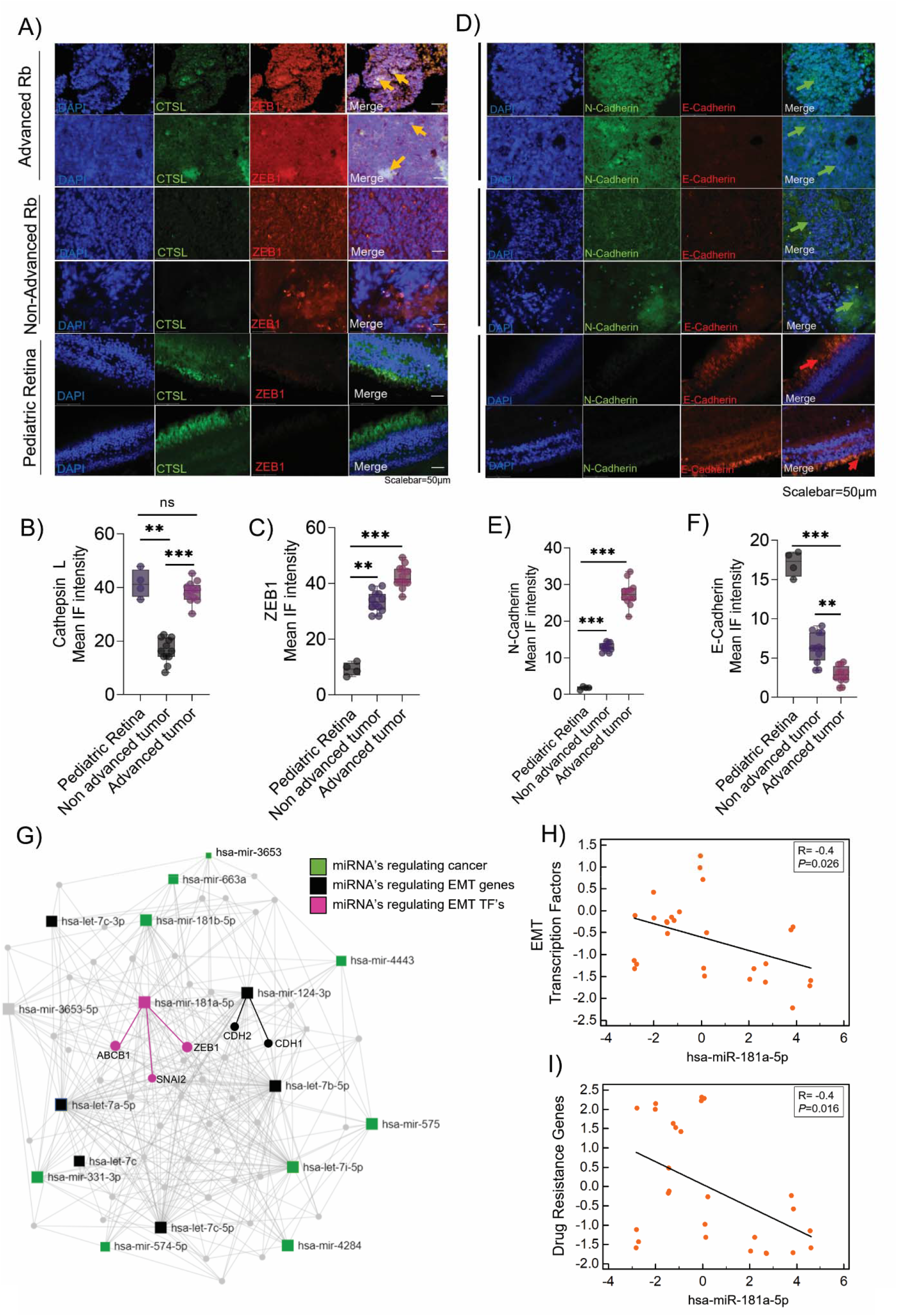
Validation of epithelial to mesenchymal transition (EMT) and chemo-drug resistance proteins in Rb tumors and their interaction with miR-181a-5p. Immunofluorescence showing expression of (A) ZEB1 and CTSL. IF mean intensity of (B) ZEB1 and (C) Cathepsin L staining in advanced (n=12), non-advanced (n=12) and control pediatric retina (n=4). Immunofluorescence showing expression of (D) N-Cadherin and E-Cadherin, IF mean intensity of (E) N-Cadherin and (F) E-Cadherin in advanced (n=12), non-advanced (n=12) and control pediatric retina tissues (n=4). Scale bar=50µm. (G) Network map showing predicted interaction of miRNA-mRNA targets using miRNet. Correlation plot showing (H) negative correlation of EMT genes (ZEB1, SNAI2, TWIST) with miR-181a-5p in Rb tumors, (I) negative correlation of drug-resistant genes (MDR1, MRP1 and CTSL) with miR-181a-5p in Rb tumors. Values represents mean ± s.d. Two tailed Mann-Whitney was used for statistical analysis. *p < 0.05, **p < 0.01, ***p < 0.001.

### Chemotherapy resistant Rb cells confer enhanced EMT and invasion

Initial regression of the Rb tumors post treatment is followed by an orbital relapse or recurrence of a more aggressive chemo-resistant tumors composed of tumor cells with a much higher tumor initiating ability than the original tumor (Cicinelli & Kaliki, 2019). EMT program through *ZEB1* is known to drive cellular mobility and tumor dissemination in other types of cancer (Drapela et al., 2020), however their role in EMT driven drug resistance in Rb tumors are unknown. To extend our study of the consequences of *RB1* downregulation in Rb tumors and its influence on miRNA and EMT signatures, we overexpressed *RB1* gene in Rb null Y79 cells. In real time gene expression assays, we found miR-181a-5p to be significantly upregulated in the presence of Rb compared to Rb null cells (*P*=0.002) (Figure 3A). Rb over expression decreased key EMT factors like ZEB1, Slug, N-cadherin and drug resistant MDR1 proteins (Figure 3B). However, Rb over expression increased E-cadherin expression indicating a halt in the EMT switch (Figure 3B). These findings strongly suggest that Rb mediates the suppression of EMT and in absence of Rb, EMT drives a mesenchymal phenotype and drug resistance to further promote invasion and migration (Figure 3C). To further elucidate the mechanism, we developed Y79 cells resistant to topotecan and carboplatin by exposing them to increasing concentrations of the drugs for 3 weeks (Figure 3C). After each week, the surviving cells that reached >60% confluency were passaged in fresh media with increased concentration of topotecan or carboplatin. The procedure was performed repeatedly until the cells display low sensitivity to IC50 doses of topotecan or carboplatin (Figure 3D), resistance to DNA damage defined by low λH2A.X foci count under an IC50 dose therapy for 48hours (Figure S2A, B, C), shift in IC50 values of topotecan and carboplatin (Figure S2D, E) and high surface expression of MDR1 proteins (Figure 3F, G, H, S2F, G, H), marking a resistant phenotype. We developed tumor spheroids for parental and resistant Y79 cells, and we observed high ZEB1 and cathepsin L expression in resistant spheroids by immunofluorescence. However, parental spheroids displayed strong ZEB1 expression and no cathepsin L expression (Figure 3I). We further confirmed the findings using RT-PCR that revealed a mesenchymal transition trend for resistant lines compared to parental lines (Figure 3, J, K, L, M, S2I). Likewise, using transwell assay, we found increased invasion and migration of resistant Y79 cells compared to sensitive Y79 cells, under high dose topotecan (100nM) therapy (Figure 2N, O, P). In line with the above results, in human Rb tumors, miR-181a-5p was significantly downregulated in EMT-high/ drug resistant advanced tumors. Our results point to miR-181a-5p as a potential negative regulator of the EMT and drug resistance mechanisms that could possibly influence tumor metastasis (Figure S2J).

**Figure 3.**
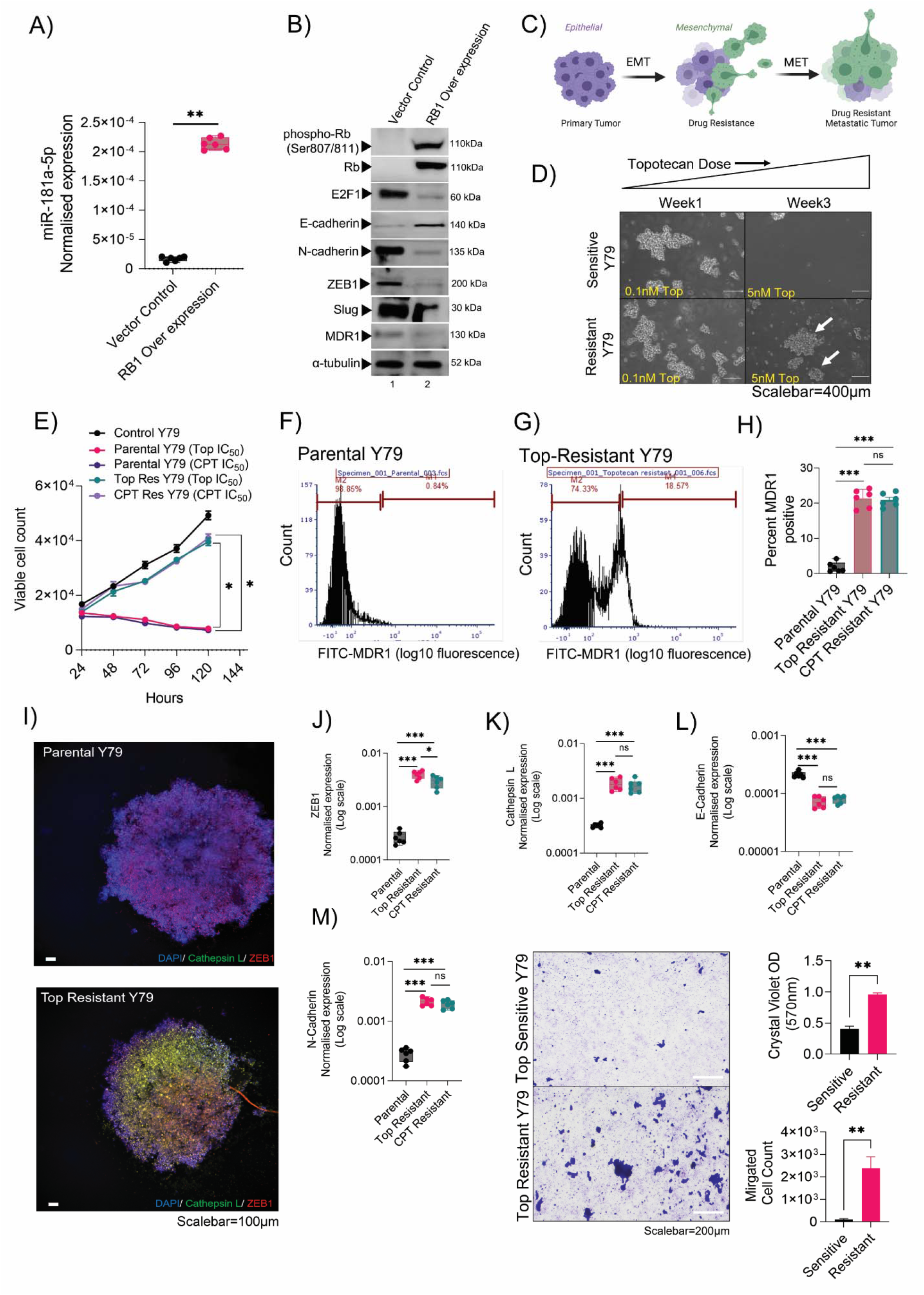
Chemotherapy resistant Rb cells confer enhanced EMT and invasion. (A) RT-PCR showing normalized expression of miR-181a-5p in Vector control (*RB1* null) and *RB1* over expressed Y79 retinoblastoma cells. (B) Immunoblot showing expression of EMT and chemoresistant markers in Vector control (*RB1* null) and *RB1* over expressed Y79 cells. (C) Schematic showing EMT program and drug resistance induction in metastatic tumors. (D) Phase contrast microscopy images showing morphology of parental and resistant Y79 cells under increasing dose of topotecan treatments from week1 to week3. Scalebar=400µm. (E) Cell viability of parental, topotecan resistant and carboplatin resistant Y79 cells at 24hr, 48hr, 72hr and 96hr. MDR1 surface expression analysis in (F) parental and (G) topotecan resistant Y79 cells by flow cytometry. (H) Bar graph showing the percentage of cells positive for MDR1 surface expression in parental, topotecan resistant and carboplatin resistant Y79 cells. (I) Parental and resistant Y79 spheroids showing expression of ZEB1 and Cathepsin L. Scale bar= 100µm. RT-PCR showing expression of (J) ZEB1 (K) Cathepsin L (L) E-cadherin and (M) N-cadherin in parental, topotecan resistant and carboplatin resistant Y79 cells. (N) Transwell invasion and migration assay to assess the migratory capacity of resistant cells compared to sensitive cells under 10nM topotecan treatment for 48hours. (O) Crystal violet OD reading at 570nm to assess invasiveness (P) Trypan blue count to assess migrated cells in the lower compartment of the transwell chamber. N=3, Two-tailed Student’s *t*-test (for 2< group) and one-way ANOVA with Dunnett’s multiple comparisons tests (for >2 group) were used for statistical analysis. *p < 0.05, **p < 0.01, ***p < 0.001.

### Resistant cells undergo transition mediated by *ZEB1* and acquire resistance through Cathepsin L

To identify the critical downstream signaling pathways that regulate EMT and chemo-resistance in the context of Rb, we evaluated the TGFβ pathway, as it was highlighted by the pathway analysis from the transcriptomic profile of advanced Rb tumors (Figure S3A). SMAD2 phosphorylation was enhanced in resistant Y79 compared to the parental (Figure 4A) line. The resistant cells showed more pronounced expression of EMT markers such as ZEB1, Slug, N-cadherin and drug resistance markers like MDR1 and Cathepsin L (Figure 4A) suggestive of advanced tumor cells with metastatic potential. In agreement to a previous report that retinoblastoma cells lack functional TGFβ receptor I and II (Horie et al., 1998) (Figure 4B, S3B), we identified the *ACVR1C* receptor (Activin A Receptor Type 1C), a member of the TGFβ family, known to activate SMAD2 in advanced retinoblastoma (Asnaghi et al., 2019) (Figure S3C). Resistant cells had increased expression of *ACVR1C*, indicating its likely role in activating SMAD2 dependent TGFβ signaling. To evaluate if TGFβ modulation affects chemoresistance, we used TGFβ ligand activation (10ng) and TGFβ inhibitor (50µM SB43152) for 48hours in parental and resistant lines and assessed the changes in EMT and drug resistance markers. We found, by immunofluorescence, enhanced levels of phospho-SMAD2 and cathepsin L levels upon TGFβ activation in resistant cells (Figure 4D), while TFGβ activated parental cells showed lesser induction of phospho-SMAD2 and Cathepsin L levels compared to controls. Likewise, TGFβ activation also increased ZEB1 expression in resistant and parental cells compared to controls (Figure S3D). TGFβ inhibition in parental lines shows complete reduction of phospho-SMAD2, ZEB1 and cathepsin L proteins, however resistant cells upon TGFβ inhibition shows partial reduction of cathepsin L and ZEB1 in the nucleus (Figure 4D, S3D). Consistently, lower levels of ZEB1, Slug and Cathepsin L proteins were observed upon TGFβ /phospho-SMAD2 inhibition in resistant lines (Figure 4E). In contrast to the resistant phenotype, TGFβ/SMAD2 inhibition drastically reduced ZEB1, slug and depleted cathepsin L proteins in parental lines. TGFβ inhibition showed significant downregulation of *SMAD2* (Figure S3E) and *ZEB1* genes in the resistant lines (Figure 4F), while TGFβ inhibition did not affect *SNAI2* and *CTSL* (Cathepsin L) expressions in resistant cells (Figure 4G, H). Thus, TGFβ transcriptionally regulates *ZEB1*, but not *SNAI2* and *CTSL* in resistant lines (Figure 4F, G, H). Using in silico promoter sequence analysis, we found that *ZEB1* promoter has direct binding sites for *SMAD2* located in close proximity to the cognate TSS (transcriptional start site), suggestive of its potential for transcriptional activation of *ZEB1* (Figure S3F). However, *SMAD2* binding sites in *SNAI2* promoter are relatively distant from the TSS (Figure S3F), a potential reason for its poor sensitivity to TGFβ inhibitors. Notably, *SNAI2* also has binding sites in the *CTSL* promoter, therefore, it is possible that the transcriptional activation of *CTSL* is mediated via *SNAI2* and not *SMAD2* or *ZEB1* (Figure S3F). In support to our results, we observed enhanced nuclear localization of cathepsin L in resistant lines indicating its transcriptional activity independent of TGFβ/SMAD2 signals (Figure S3G). We speculate that resistant cells have nuclear localization of *CTSL* due to the lack of cystatin B (*CSTB*), that is known to inhibits *CTSL* nuclear localization (Ceru et al., 2010) and low expression of *CTSB* was also evidenced in the advanced tumor microarray profile. This indicates that *ZEB1* triggers EMT through TGFβ and the activated EMT program through *SNAI2* regulates *CTSL* mediated chemo resistance in advanced Rb tumors (Figure 4I). Hence, identifying a common regulator like miR-181a-5p that governs the mechanisms of both transition and resistance is both reasonable and promising for effective management of metastatic dissemination.

**Figure 4.**
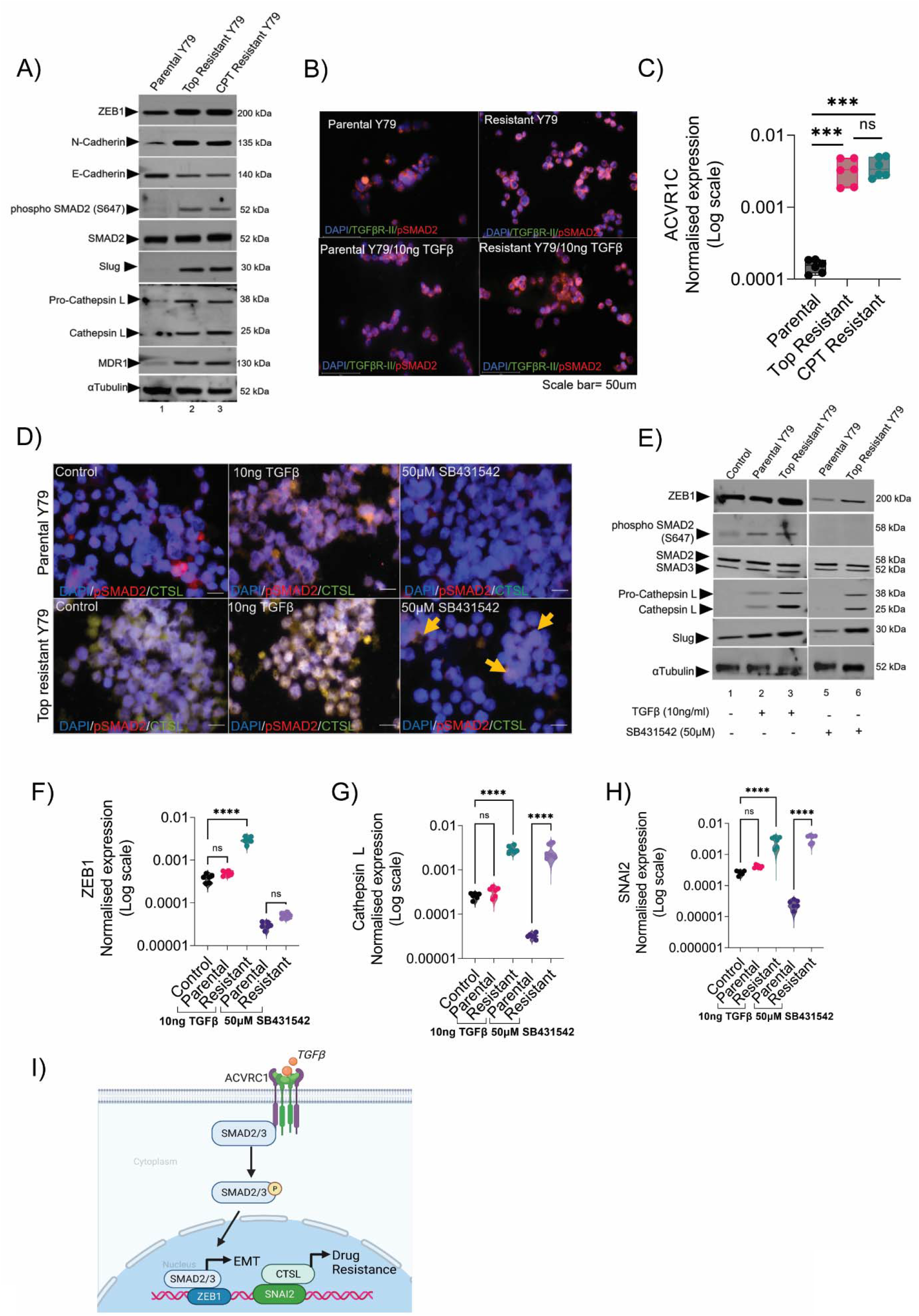
Resistant cells undergo transition mediated by *ZEB1* and acquire resistance through Cathepsin L. (A) Immunoblot showing the expression of EMT and drug resistance markers in parental, topotecan resistant and carboplatin resistant Y79 cells. (B) Immunofluorescence showing expression of TGFβ R-II and phospho-SMAD2 in parental and topotecan resistant Y79 cells with and without TGFβ induction. Scale bar=50µm. (C) RT-PCR results showing expression of ACVR1C in parental, topotecan resistant and carboplatin resistant cells. (D) Immunofluorescence showing expression of phospho-SMAD2 and cathepsin L (CTSL) upon TGFβ induction (10ng for 48 hours) and TGFβ inhibition (50µM SB431542 for 48 hours) in parental and topotecan resistant Y79 cells. Scalebar=50µm. (E) Immunoblot showing EMT and drug resistance pathway protein levels upon TGFβ induction and inhibition for 48 hours. RT-PCR results showing normalized expression of (F) *ZEB1* (G*) SNAI2* (H) *CTSL* (Cathepsin L) upon TGFβ induction and inhibition for 48hours. (I) Schematic showing the novel regulation of EMT and drug resistance mechanism in Rb tumors. N=3, Two-tailed Student’s *t*-test (for 2< group) and one-way ANOVA with Dunnett’s multiple comparisons tests (for >2 group) were used for statistical analysis. *p < 0.05, **p < 0.01, ***p < 0.001, ****p < 0.0001

### Augmenting miR-181a-5p levels confers sensitivity to chemotherapy

To evaluate if miR-181a-5p can directly regulate EMT and chemoresistance in resistant Y79 lines, we measured ZEB1, Slug and Cathepsin L proteins. Using bioinformatic tools, we also predicted the binding sites of miR-181a-5p in *ZEB1* and *SNAI2* 3’ UTR regions (Figure S3 H, I). In contrast to a non-targeting mimic control, transfection of miR-181a-5p drastically reduced ZEB1, Slug and Cathepsin L protein levels (Figure 5A). Conversely, miR-181a-5p inhibition shows increased expression of ZEB1, Slug and Cathepsin L, mimicking an EMT high, drug resistant advanced tumor phenotype (Figure 5A). Cell surface expression of MDR1 was altered in resistant cells complemented with miR-181a-5p (Figure 5B, C), that were partly explained by changes in levels of Slug and Cathepsin L. Notably, augmenting miR-181a-5p reduced cell proliferation (Figure 5D), invasion and migration of resistant cells (Figure 4E, F, G), while its inhibition showed opposite results in resistant lines (Figure 5D, E, F, G). These findings suggest that enhancing miR-181a-5p in resistant lines could sensitize them to low dose chemotherapy. We compared the response of miR-181a-5p modulated resistant lines to IC50 dose of topotecan (parental Y79_IC50_=10nM) for 96 hours. Following topotecan treatment, miR-181a-5p mimic transfected resistant lines show no significant response at 24hour and 48hour (Figure 5H, I), while they showed an increased sensitivity and low survival to topotecan_IC50_ by 72 hours (Figure J) and 96 hours (Figure K). Together, the results suggest that the miR-181a-5p plays a major role in the depletion of EMT and drug resistance phenotype by sensitizing the cells to low dose chemotherapy.

**Figure 5.**
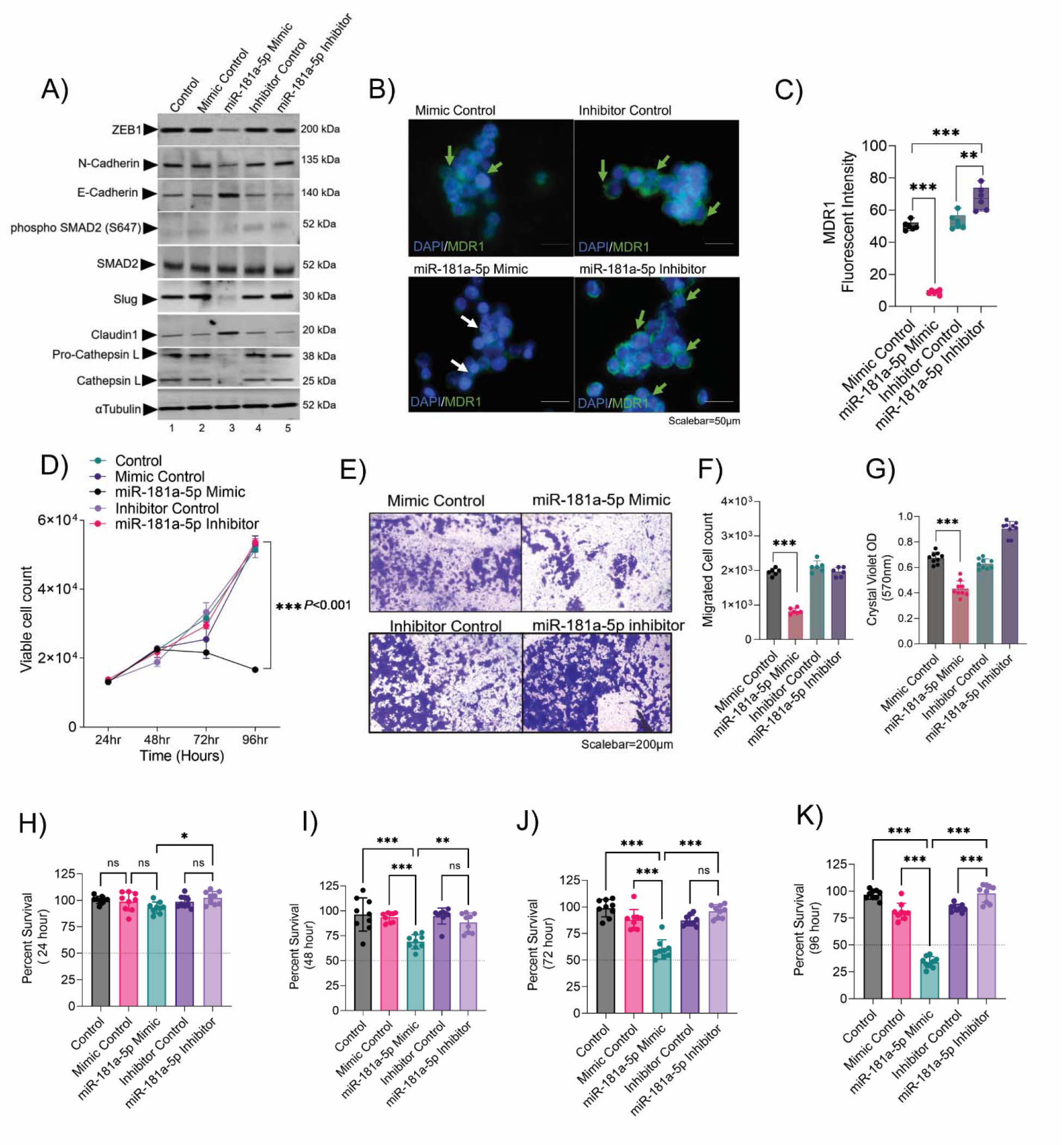
Augmenting miR-181a-5p levels confers sensitivity to chemotherapy. (A) Immunoblot showing expression of key EMT factors and drug resistance markers upon miR-181a-5p overexpression and inhibition in topotecan resistant Y79 cells. (B) Immunofluorescence showing MDR1 surface expression in topotecan resistant Y79 cells upon miR-181a-5p overexpression and inhibition. Scalebar=50µm. (C) Bar graphs showing MDR1 fluorescent intensity in topotecan resistant Y79 cells upon miR-181a-5p overexpression and inhibition. (D) Trypan blue cell count showing proliferation of topotecan resistant cells at 24hr, 48hr, 72hr and 96hr upon miR-181a-5p overexpression and inhibition. (F) Transwell invasion and migration assay to assess the invasive and migratory capacity of topotecan resistant cells upon miR-181a-5p overexpression and inhibition (F) Crystal violet OD measurement at 570nm to assess the invasiveness of resistant Y79 cells. (G) Trypan blue cell count shows migrated cells in the lower compartment of the transwell chamber. Chemosensitivity of miR-181a-5p modulated topotecan resistant Y79 cells upon 10nM topotecan treatment for (H) 24 hours (I) 48hours (J) 72 hours (K) 96 hours. Control represents untreated topotecan-resistant Y79 cells. N=3, Two-tailed Student’s *t*-test (for 2< group) and one-way ANOVA with Dunnett’s multiple comparisons tests (for >2 group) were used for statistical analysis. *p < 0.05, **p < 0.01, ***p < 0.001.

## Discussion

The present study identifies miR-181a-5p as a previously unrecognized regulator of EMT transcription factors and chemotherapy resistance. While the study focuses on intraocular advanced and non-advanced retinoblastoma tumors, our findings can be extended to other cancer systems that have persistent EMT associated chemotherapy resistance. We found miR-181a-5p to be significantly downregulated in advanced Rb tumors and provide evidence that mesenchymal transition and chemoresistance in tumors are likely sensitive to chemotherapy when miR-181a-5p is complemented.

Context dependent tumor promoting (Xue et al., 2020; Yang et al., 2018) and tumor suppressing (Li et al., 2015; Liu et al., 2020) roles have been reported for miR-181-5p. However, we report the downregulation of miR-181a-5p in retinoblastoma tumor tissues from patients when compared to healthy pediatric retina. It should be noted that the significantly reduced miR-181-5p levels in tumors observed in our study is likely due to the use of pediatric healthy retina as controls. The miRNA 181a-5p also plays a functional role in retina (Chen et al., 2020), hence the consistent low expression of miR-181a-5p expression in advanced and non-advanced retinal tumors is important in the context of the dedifferentiated tumor tissue.

The control of EMT and chemotherapy resistance by miR-181a-5p evidenced here could provide an explanation for the apparent complex roles of miR-181a-5p in advanced tumors. Recent evidence indicates that EMT occurs through intermediate states rather than being a binary process (Pastushenko et al., 2018) and is partially reactivated in various cancers (Zhang & Weinberg, 2018). We propose that dynamic reduction in miR-181a-5p levels in tumor, particularly between the different stages of Rb progression may contribute to the context dependent EMT plasticity from cancer initiation to metastasis. Differences in EMT and drug resistance transcripts between control and tumor tissues and different stages of Rb further contribute to this complexity, as miR-181a-5p may control EMT in a tissue and function specific manner. In this study, advanced Rb tumors showed increased expression of EMT signatures, a consequence of miR-181a-5p downregulation, as an EMT trigger. We propose that the EMT genes like *ZEB1* and *SNAI2* acquires transcript stability in advanced tumors, at least partly, due to reduced miR-181a-5p based degradation (Baek et al., 2008). Furthermore, we have identified chemotherapy resistance pathway genes like *ABCB1* (MDR1) (Krech et al., 2012) and *CTSL* (Cathepsin L)(Zheng et al., 2009) as EMT targets in Rb tumors. Likewise, the miR-181a-5p clusters located on chromosome 1 are known to repress E2F transcription factors (Lin et al., 2018), G1/S cell cycle regulators (Shen et al., 2018) and proto-oncogenes (Ouyang et al., 2022). We have found that miR-181a-5p negatively controls EMT and chemoresistance through regulation of *SNAI2* and *CTSL* transcripts invitro. Collectively, our work reveals an emerging and intriguing feature of miR-181a-5p and its association with a variety of distinct signaling pathways in Rb tumors.

We identified the balance between EMT driven metastasis in Rb tumors to be influenced by TGFβ in the tumor microenvironment, which is a known promoter of EMT in Rb depleted tumors (Joseph et al., 2014). Previous studies have highlighted the lack of canonical TGFβ receptors in Rb cells (Horie et al., 1998) and in agreement with recent reports (Asnaghi et al., 2019), we show that TGFβ signals acts through *ACVR1C* receptors and activates SMAD2/3 effectors in Rb. Mechanistically, we show that the presence of TGFβ ligand mediates a mesenchymal shift in Rb cells and is associated with enhanced migration and invasion capacity. Conversely, treatment with TGFβ signaling inhibitor reduced *ZEB1, SNAI2* levels and prevented mesenchymal marker expression and morphological changes, thus linking mesenchymal differentiation in Rb with enhanced tumor cell invasion through the TGFβ/ZEB1/SNAI2 axis. Interestingly, in line with our observations regarding miRNA mediated regulation of EMT and chemoresistance, a recent report highlights that the miR-200c-ZEB1 feedback loop is involved in invasion, migration and chemoresistance in advanced glioblastoma tumors (Siebzehnrubl et al., 2013). We have found that TGFβ signals can drive an EMT program in Rb-/- cells, while they do not necessarily lead to chemoresistance. We observed that Rb cells acquire chemotherapy resistance through enhanced EMT program orchestrated by *SNAI2* by regulating *CTSL.* This concept is further supported by our observations with ectopic expression of miR-181a-5p, that repress *SNAI2* at its 3’ UTR region and targets the SNAI2-CTSL signaling cascade, inhibiting transition and resistance.

The tumor suppressor properties of miRNA in various cancers have prompted the development of various potent inhibitors of pharmacological targeting in clinical settings (Bonci et al., 2008). Although our study was limited to *in vitro* models, our findings on miR-181a-5p raises hopes for therapeutic strategies for the management of advanced Rb tumors. In conclusion, our work reveals a unique mechanistic link between EMT and chemoresistance in Rb tumors, illustrating miR-181a-5p as a potential therapeutic target.

## Materials & Methods

### Clinical samples

The study was conducted in accordance to the principles outlined in the Declaration of Helsinki under a protocol approved by the institutional ethics committee of Narayana Nethralaya (EC Ref no: C/2013/03/02). Informed written consents were received from all parents before inclusion in the study. Histology confirmed Rb tumors (n=9) comprising of Group E and Group D of the age range 0.2 - 4years and pediatric controls (n=2) of the age range (0.2-0.3 years) were used for the miRNA and mRNA microarray study. The details of clinical samples including age, gender, laterality, tumor viability, clinical and histopathology details are mentioned in Table 1. For immunohistochemistry validations, we have used additional subset of Rb subjects comprising of Group E (n=4) and Group D (n=4) and pediatric retina (n=4) of the age range 0.2-4 years. The details of clinical and histopathology details of additional Rb subjects are mentioned in Table S1.

### Tumor miRNA and mRNA profiling

Total RNA was isolated from 9 Rb tumors and 2 control pediatric retina samples using Agilent Absolutely RNA miRNA kit (cat# 400814) according to the manufacturer’s instructions. The quality of isolated RNA was determined on an Agilent 2200 TapeStation system (G2964AA) using an Agilent RNA ScreenTape assay (5067-5576). mRNA labeling and microarray processing was performed as detailed in the “One-Color Microarray-Based Gene Expression Analysis” (v 6.9, cat# G4140-90040). miRNA labeling was done using an Agilent miRNA Complete Labeling and Hyb Kit (Cat# 5190-0456). The gene expression and miRNA data were extracted using Agilent Feature Extraction Software (11.5.1.1) and analyzed using Agilent GeneSpring GX 13.1. Statistical analysis was carried out using a t-test unpaired statistical method with Benjamini Hochberg FDR method. In both mRNA and miRNA analyses, transcripts exhibiting P ≤ 0.05 and fold changes greater than or equal to two were differentially expressed.

### Cell lines

Y79 cells were obtained from American Type Culture Collection (ATCC, Manassas, VA). The Y79 cells were cultured in RPMI 1640 medium (Gibco, Cat #11875093) supplemented with 10% FBS and 1% Pen Strep (Penicillin –Streptomycin) and maintained at 37°C in a humidified atmosphere of 5% CO_2,_ with intermittent shaking in an upright T25 flask. To generate chemotherapy-resistant lines, Y79 cells were exposed to media containing a low dose (1/100^th^ of IC50) of topotecan or carboplatin for 48hours and replenished with fresh media without drugs for the next 48hours and vice versa. At the end of each week, we increased the dose of topotecan and carboplatin by 10folds for 3-4 weeks, till the cells display tight large clusters and no sensitivity to chemo-drugs. The cells were further analysed for MDR1 surface expression and IC50 shift to confirm the resistant phenotype.

### Gene expression analysis

Total RNA extracted from the second cohort of clinical subjects was used for RT PCR validation for mRNA and miRNA microarray. RT-PCR was performed with Agilent Brilliant III Ultra-Fast RT-PCR reagent (cat# 600884), using Agilent AriaMX real-time PCR instruments. Relative mRNA expression was quantified using the ΔΔC(t) method (Livak & Schmittgen, 2001). For in-vitro assays, total RNA was isolated from cells using the Trizol reagent (Invitrogen, Carlsbad, CA) according to the manufacturer’s protocol. 1µg of RNA was reverse transcribed using Bio-Rad iScript cDNA synthesis kit (cat# 1708890) and quantitative real-time PCR was performed using Kappa Sybr Fast qPCR kit (cat# KK4601) using Bio-rad CFX96 system. Relative mRNA expression levels were quantified using the ΔΔC(t) method. Results were normalized to housekeeping human β-actin. Details of primers used are described in the Table 2 below.

**Table 2:**
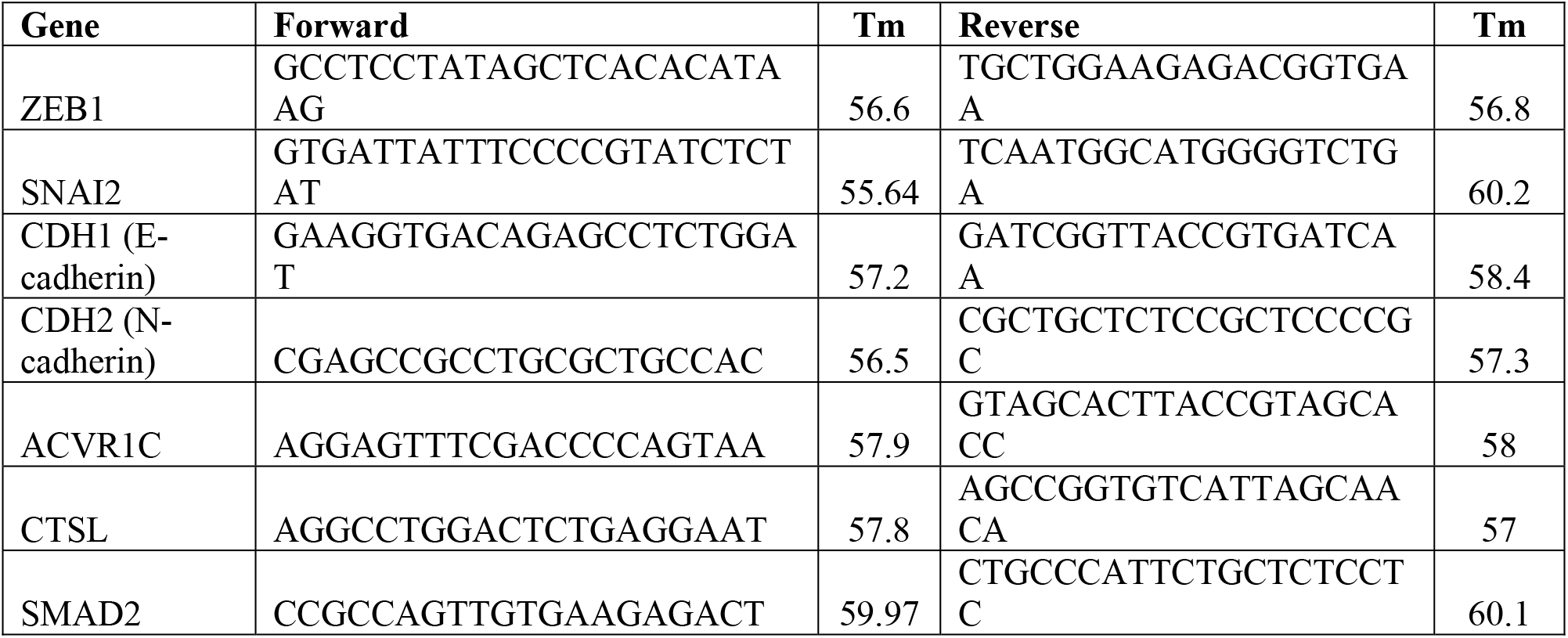
Details of qPCR primers used in the study.

For qPCR of miRNAs, miRNA was converted to cDNA using the reverse transcription kit, miRCURY LNA Universal RT microRNA PCR (cat#339306, Qiagen). Briefly, RNA was polyadenylated with ATP by poly(A) polymerase at 37°C for 1 hr and reverse transcribed using 0.5 μg of poly(T) adapter primer. Each miRNA was detected by the mature DNA sequence as the forward primer and a 3′ universal reverse primer provided in the QuantiMir RT kit. Human small nuclear U6 RNA was amplified as an internal control. qPCR was performed using Power SYBR Green PCR Master Mix (Applied Biosystems). All qPCR performed using SYBR Green was conducted at 50°C for 2 min, 95°C for 10 min, and then 45 cycles of 95°C for 10 s and 60°C for 1 min. The specificity of the reaction was verified by melt curve analysis. The details of miRNA primers used are mentioned in Table 3.

**Table 3:**
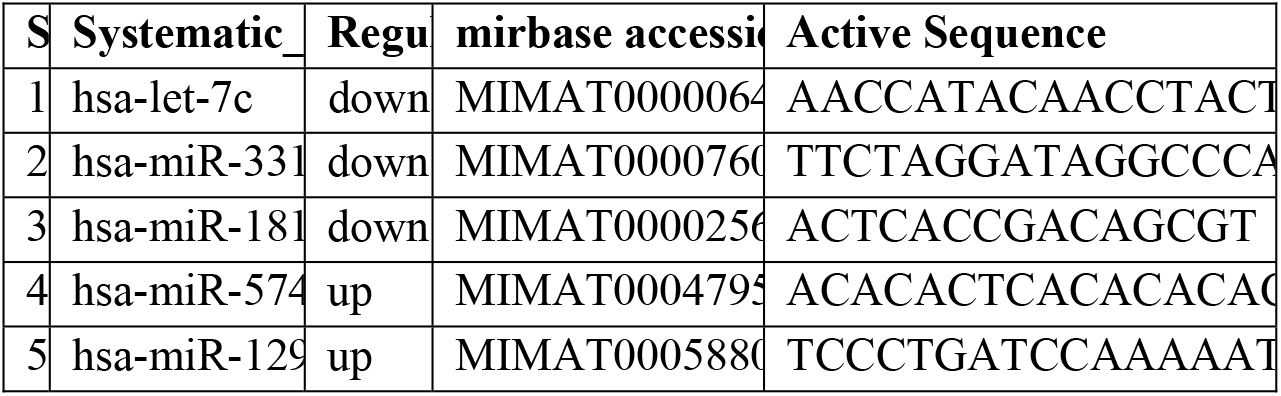
Details of miRNA qPCR primers.

### Histopathology & light microscopy

Paraffin-embedded specimens of Rb tumor and control retina were used. 4µm paraffin sections were dewaxed at 60°C, rehydrated in decreasing concentration of ethanol. Slides were stained with hematoxylin & eosin according to standard procedures. Brightfield images were captured using Olympus CKX53 microscope.

### Immunofluorescence

For IF, 4µm sections of Rb tumor and pediatric retina were deparaffinized, rehydrated & were subjected to heat-induced epitope retrieval using citrate buffer for 20 min at 100 °C. After 2% BSA block, tissues were incubated overnight at 4°C with antibodies for ZEB1 (1:1000; cat#70512, Cell Signaling Technology), Cathepsin L (1:500; cat#ab6314, Abcam), E-cadherin (1:500; cat#3195, Cell Signaling Technology), N-cadherin (1:1000; cat#14215, Cell Signaling Technology), MDR1 (1:1000, cat#13342, Cell Signaling Technology). For in-vitro experiments, 2×10^3^ parental and topotecan resistant Y79 cells were seeded on 8-chamber glass slides, precoated with poly-L-lysine. The cells were stained with phospho λ-H2A.x (ser139) (1:500; cat#9718, Cell Signaling Technology), ZEB1 (1:500; cat#70512, Cell Signaling Technology), Cathepsin L (1:500; cat#ab6314, Abcam), MDR1 (1:500, cat#13342, Cell Signaling Technology) phospho-SMAD2 (1:1000, cat#ab53100, Abcam) and TGFBR2 (1:1000, cat#ab78419, Abcam). Secondary antibodies used include goat anti-mouse Alexa Fluor 488 (1:5000, cat# ab150113, Abcam) and donkey anti-rabbit Cy3 (1:5000, cat# 711-1650152, Jackson ImmunoResearch Laboratories). Hoechst 33342 (1:5000) was used for nuclear staining. Images were analyzed and captured using EVOS M7000 imaging systems (ThermoFisher Scientific). The fluorescent intensity was measured using ImageJ software (NIH Image, Bethesda, MD).

### Western blotting

For Western blot analysis, cells were lysed in RIPA buffer (20mM Tris pH 8.0, 0.1% SDS, 150 mM NaCl, 0.08% Sodium Deoxycholate, 1% NP40 supplemented with 1 tablet of protease inhibitor (Complete ultra mini-tablet, Roche) and phosphatase inhibitor (PhosphoStop tablet, Roche). 20µg of total protein was loaded per lane and were separated by SDS-PAGE. The separated proteins on the gel were transferred onto PVDF membrane and were probed for specific antibodies against Rb (cat# 9309; Cell signaling) phospho-Rb (cat# 8516, Cell signaling), E2F1 (1:500, cat#sc-251, SantaCruz Biotechnology), ZEB1 (1:1000; cat#3396, Cell Signaling Technology), Cathepsin L (1:1000; cat#ab6314, Abcam), Slug (1:1000; cat#9585, Cell Signaling Technology), E-cadherin (1:1000; cat#3195, Cell Signaling Technology), N-cadherin (1:1000; cat#14215, Cell Signaling Technology), MDR1 (1:1000, cat#13342, Cell Signaling Technology), Total smad2/3 (1:1000, cat#ab207447, Abcam), phospho-SMAD2 (1:1000, cat#ab53100, Abcam), TGFBR1 (1:1000, cat#PA1731, Boster Bio), TGFBR2 (1:1000, cat#ab78419, Abcam), α-Tubulin (1:1000, cat# 3873; Cell signaling) and GAPDH (1:1000, cat#5174; Cell signaling) in 5%BSA in 1xTBST, overnight at 4°C. For nuclear-cytoplasmic fractionation, the cytoplasmic fraction was extracted using a hypotonic buffer for 30min on ice and the nuclear fraction was extracted using a lysis buffer solution containing 10 mM Tris at pH 8, 170 mM NaCl, 0.5% NP40 with protease inhibitors. The respective cellular fractions were incubated with respective primary antibodies for immunoprecipitations. LaminA/C (1:1000, sc-6215, SantaCruz Biotechnology) was used as a nuclear fraction loading control and α-Tubulin as a cytoplasmic fraction loading control (1:1000, cat# 3873; Cell signaling).

After 4 washes with 1x TBST for 10 minutes, membranes were incubated HRP-conjugated anti-mouse (cat#7076; Cell signaling) or anti-rabbit antibodies (cat#7074; Cell signaling) at 1:2000 dilution for 2 h. Images were visualized using the Image Quant LAS 500 system (GE Healthcare Life Sciences, USA).

### FACS analysis of MDR1 surface staining

The cells surface expression of MDR1 in parental and resistant Y79 cells was detected using an anti-MDR1 antibody (1:500, cat#13342, Cell Signaling Technology). Parental and resistant Y79 cells post-drug exposure was incubated in 200µl of PBS containing 1% FBS and 2ug of MDR1 antibody at 4°C for 1hour in an intermittent shaker. After three washes with ice-cold PBS, the cells were further incubated in goat anti-rabbit Alexa Fluor 488 secondary antibody for 30minutes at RT. The cells were then washed in ice-cold PBS and analyzed with FACS apparatus equipped with FACSDiva software. The fluorescent intensity of the FL1 channel was plotted to compare the cell surface expression of MDR1 in parental and resistant lines.

### Cell proliferation assay

Parental Y79, Topotecan resistant Y79, and Carboplatin resistant Y79 cells were used for the proliferation assay. 10000 Y79 cells were seeded in 24 well plates for proliferation assay. Cell viability was determined once every 24hours for 4 consecutive days using trypan blue cell staining and cell counting using a hemocytometer. In miRNA transfected models, 10000 cells were seeded onto 24 well plates post 48hours of transfection, and proliferation was assessed from 24 hours to 96hours. The cell viability was determined using a trypan blue assay. The experiments were performed in three experimental repeats in triplicates for different experimental conditions. Data were expressed as mean ± SD of triplicate experiments.

### Cell migration & invasion assays

Cell migration & invasion assays were performed in 24-well transwell plates with cell culture inserts (BD Falcon). 15000 parental and resistant Y79 cells in 150μl 0% RPMI media were seeded in transwell insert coated with 1% matrigel & incubated for 48 hours. The bottom chamber was filled with 600μl of 10% RPMI media. After 48-hour incubation, cells on the insert were removed using a cotton swab. Migrated cells on the lower surface of the insert membrane were fixed with 4% PFA and stained with 0.1% crystal violet. Images were captured at brightfield using Olympus CKX53 microscope. Cells were further lysed using 10% SDS and absorbance of crystal violet was measured at 595 nm using a microplate reader. For migration assay, the cells that migrated to the bottom chamber at 48hours were counted using trypan blue cell staining and cell counting using a hemocytometer.

For miRNA transfection experiments 15000 topotecan resistant Y79 cells were seeded in 0% RPMI media in the transwell insert coated with 1% matrigel for 48hours. Invasive and migrated cells were quantified using a 0.1% crystal violet staining protocol. Data were expressed as replicate data points ± SD of triplicate experiments.

### Colony formation/ Tumor spheroid assay

The spheroid formation assays were carried out on a low attachment U-bottom 96 well plate (BRAND® 96-well microplate, Sigma Aldrich). Single-cell suspension of 500 parental and topotecan resistant Y79 cells in 10% RPMI medium was loaded in each well of a 96 well plate followed by centrifugation for 1000rpm for 1 min to facilitate cell aggregation. The cells were cultured at 37°C in a 90% humidified incubator with 5% CO2 for 7 days for the generation of tight and regular tumor spheroids. Spheroids were imaged using the EVOS FL imaging system, Invitrogen. ImageJ 2.1 software was used for spheroid area measurements. Data were expressed as replicate data points ± SD of triplicate experiments.

### Chemosensitivity assay

Cell viability of topotecan resistant Y79 cells post miRNA transfections and exposure to topotecan IC50 (10nM) treatment for 48hours was determined by Presto Blue cell viability reagent (Invitrogen) as per manufactures protocol. In brief, topotecan-resistant Y79 (5×103) were plated into 96-well plates (Eppendorf, Sigma Aldrich) and incubated overnight. Cells were treated with topotecan IC50 for 48 hours. Untreated Y79 resistant cells were considered as control. Four hours before the end of treatment, presto-blue reagent (Invitrogen) was added and incubated for 2 hours followed by measurement of fluorescence (540 nm excitation/590 nm emissions). The chemo-sensitivity of all treated cells was determined across conditions and compared against control mock-treated cells (considered as 100% viable). Data were expressed as mean ± SD of triplicate experiments.

### Pathway analysis

The KEGG functional enrichment of the gene expression microarray and miRNA microarray was carried out using the GeneSpring GX 13.1, NetworkAnalyst 3.0, and miRnet 2.0 packages; and the pathways with p < 0.05 and fold-change > 2.0 were considered significantly enriched. The construction of gene-miRNA interactions was performed and interactions network were constructed using miRnet 2.0.

### Statistical analysis

Statistical analysis was performed using GraphPad Prism 8. Data are presented as mean ± s.d unless indicated otherwise, and P < 0.05 was considered statistically significant. For all representative images, results were reproduced at least three times in independent experiments. For all quantitative data, the statistical test used is indicated in the legends. A statistical ‘decision tree’ is provided as Figure S4. Heatmaps of the Z transformed gene expression level of mRNA microarray were created using Python 3.7 Seaborne 0.11.0. Bubble weighed plots with calculated q-values were created using Python 3.6.2, circlize library.

## Author contributions

VSB and AG designed the experiments and wrote the manuscript. VSB performed cell line experiments. AM, GD and RS provided human tissue samples and helped interpret correlations with clinical data. DSA, AB, RK and NG performed the omics experiments, analyses and assisted with figure preparation and wrote relevant methods. SH edited the manuscript, helped with data analysis and scientific guidance.

## Acknowledgements

The authors thank Dr G Kumarmanickavel, Dr Swaminathan Sethu and Archana Padmanabhan Nair for their expertise and assistance throughout all aspects of our study. The authors thank Narayana Nethralaya Foundation for funding research support to VSB, AB, RK, AG. The funders had no role in design, data collection, analysis, decision to publish, or preparation of the manuscript.

## Additional information

Supplementary data figures and tables are available for this paper.

## Data availability statement

Gene and miRNA expression datasets are deposited in Gene Expression Omnibus (GEO) and their accession numbers are; Gene expression microarray (GSE208143) and miRNA expression microarray (GSE208677). All data generated or analysed during this study are included in the manuscript and supporting supplementary files.

## Supplementary Figure S1-S4

**Figure S1.**
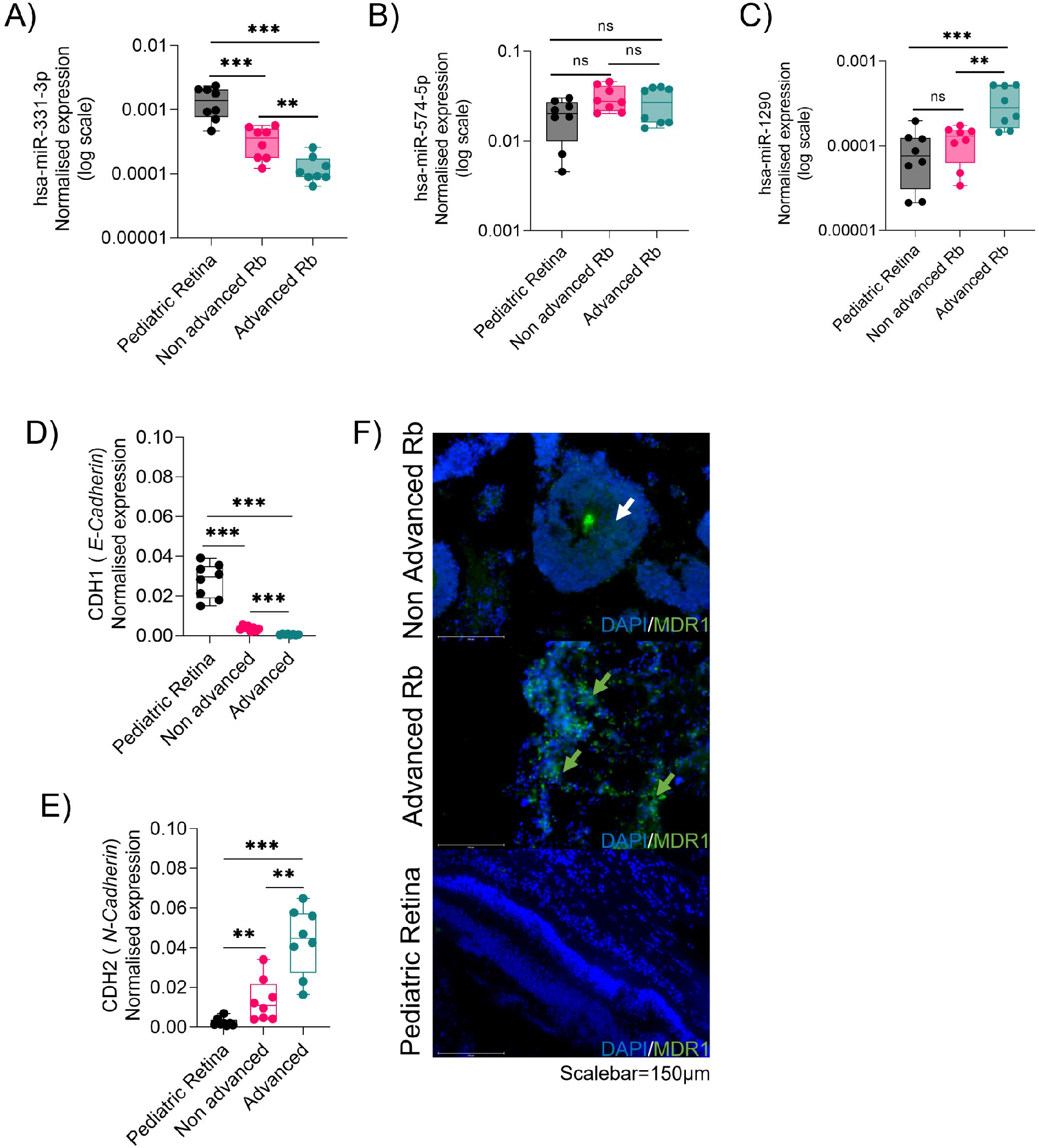
Transcriptomic profiling identifies differentially regulated miRNA’s, EMT, and drug-resistant genes in Rb tumor subtypes. RT-PCR validations of microarray identified miRNAs (A) has-miR-331-3p (B) has-miR-574-5p (C) has-miR-1290 in advanced (n=4), non-advanced (n=4) and control pediatric retina (n=4). RT-PCR validations for microarray identified mRNAs (D) CDH1 and (E) CDH2 in advanced (n=4), non-advanced (n=4) and control pediatric retina (n=4) (F) Immunofluorescence showing MDR1 expression in advanced Rb (n=4), non-advanced Rb (n=4) and pediatric retina tissues (n=4). Scale bar =150µm. Values represents mean ± s.d. Two tailed Mann-Whitney was used for statistical analysis. *p < 0.05, **p < 0.01, ***p < 0.001, ****p < 0.0001.

**Figure S2.**
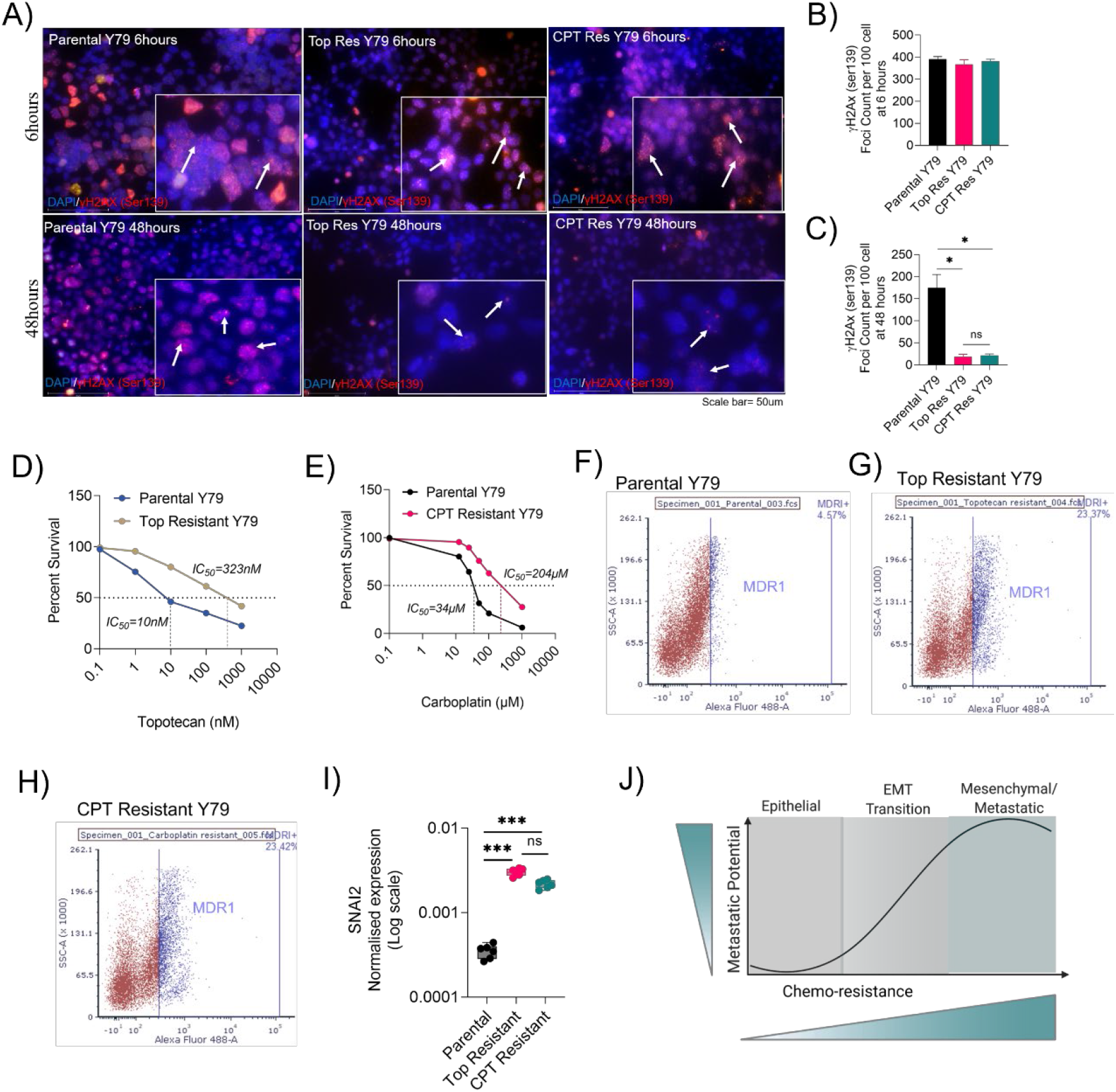
Chemotherapy resistant Rb cells confer enhanced EMT and invasion. (A) Immunofluorescence showing λH2A.x foci in parental, topotecan resistant and carboplatin resistant cells upon IC50 dose treatment using topotecan or carboplatin for 6 hours to 48hours. λH2A.x foci count at (B) 6 hours and (C) 48hours of topotecan and carboplatin therapy. Survival assay to determine the IC50 shift in resistant lines with increasing concentration of (D) topotecan (E) carboplatin. MDR1 surface staining analysed by flow cytometry in (F) Parental Y79 cells (G) Topotecan resistant Y79 cells (H) Carboplatin resistant Y79 cells. (I) RT-PCT results showing expression of SNAI2 in parental, topotecan resistant and carboplatin resistant cells. (J) Schematic showing EMT trans-differentiation and induction of drug resistance transit the cells to a dedifferentiated mesenchymal/ drug resistant metastatic phenotype. Two-tailed Student’s *t*-test (for 2< group) and one-way ANOVA with Dunnett’s multiple comparisons tests (for >2 group) were used for statistical analysis. *p < 0.05, **p < 0.01, ***p < 0.001.

**Figure S3.**
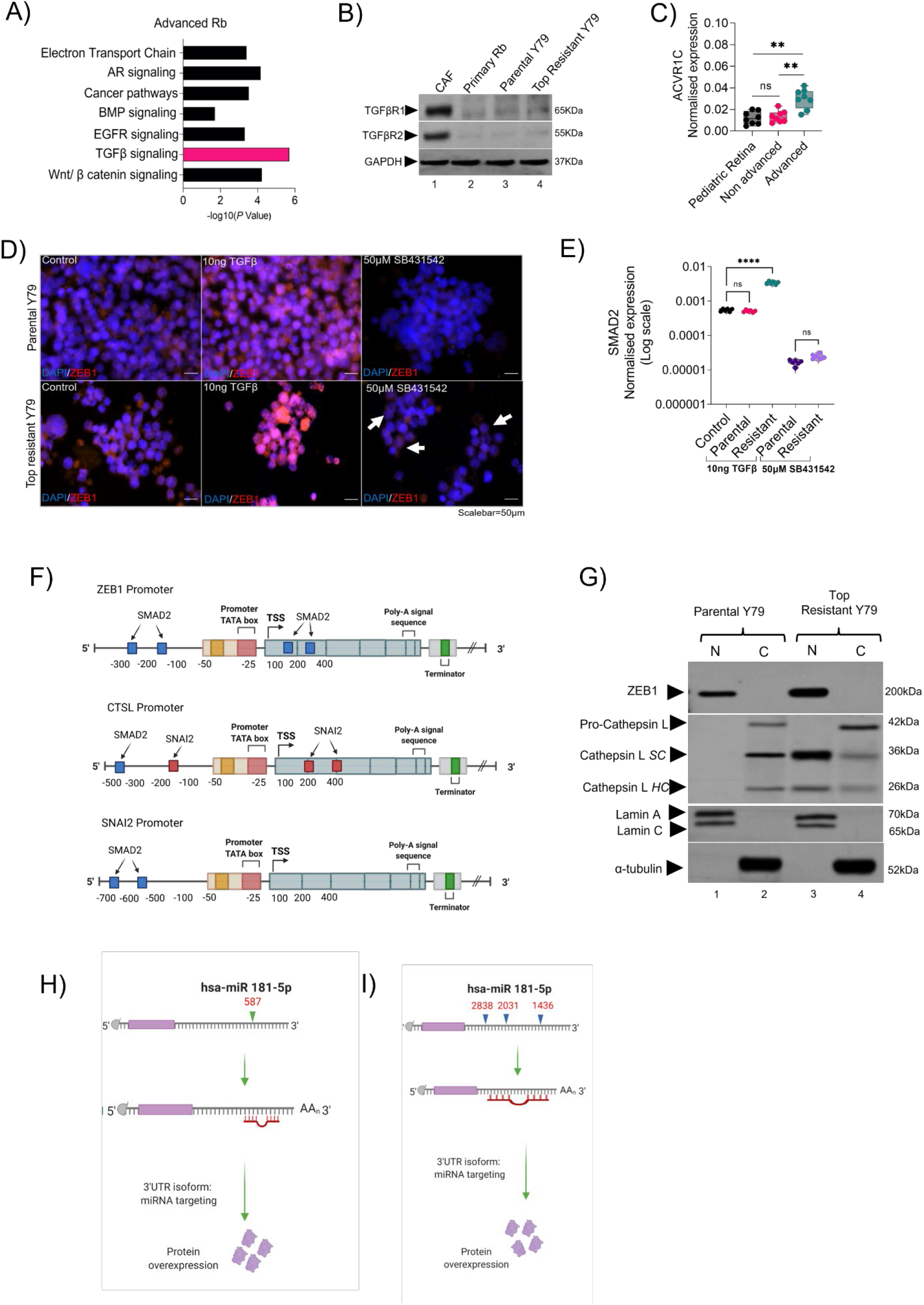
Resistant cells undergo transition mediated by *ZEB1* and acquire resistance through Cathepsin L. (A) KEGG pathway enrichment analysis showing differentially regulated pathways in advanced Rb tumors. (B) Immunoblot showing the expression of canonical TGFβ receptors I and II in retinoblastoma associated fibroblast (CAF) primary culture, T4a stage Rb tumor primary culture, parental Y79 and topotecan resistant Y79. (C) RT-PCR results showing normalized expression of ACVR1C receptors in pediatric retina (n=4), advanced (n=4) and non-advanced Rb tumors(n=4). (D) Immunofluorescence showing ZEB1 expression upon TGFβ induction and TGFβ inhibition in parental and topotecan resistant Y79 cells for 48hours. Scalebar=50µm. (E) RT-PCR showing normalized expression of SMAD2 upon TGFβ induction and TGFβ inhibition in parental and topotecan resistant Y79 cells for 48hours. (F) Schematic showing promoter binding regions of SMAD2 in ZEB1 promoter, SMAD2 and SNAI2 in CTSL promoter and SMAD2 in SNAI2 promoter. The binding sites in each promoter were curated using eukaryotic promoter database. (G) Nuclear-cytoplasmic fraction immunoblot showing the subcellular localization of ZEB1 and CTSL in parental and resistant Y79 cells. MicroRNA target prediction database (miRDB) predicted binding regions of miR-181a-5p in (H) ZEB1 3’UTR (I) SNAI2 3’UTR. Two-tailed Student’s *t*-test (for 2< group) and one-way ANOVA with Dunnett’s multiple comparisons tests (for >2 group) were used for statistical analysis. *p < 0.05, **p < 0.01, ***p < 0.001, ****p < 0.0001.

**Figure S4:**
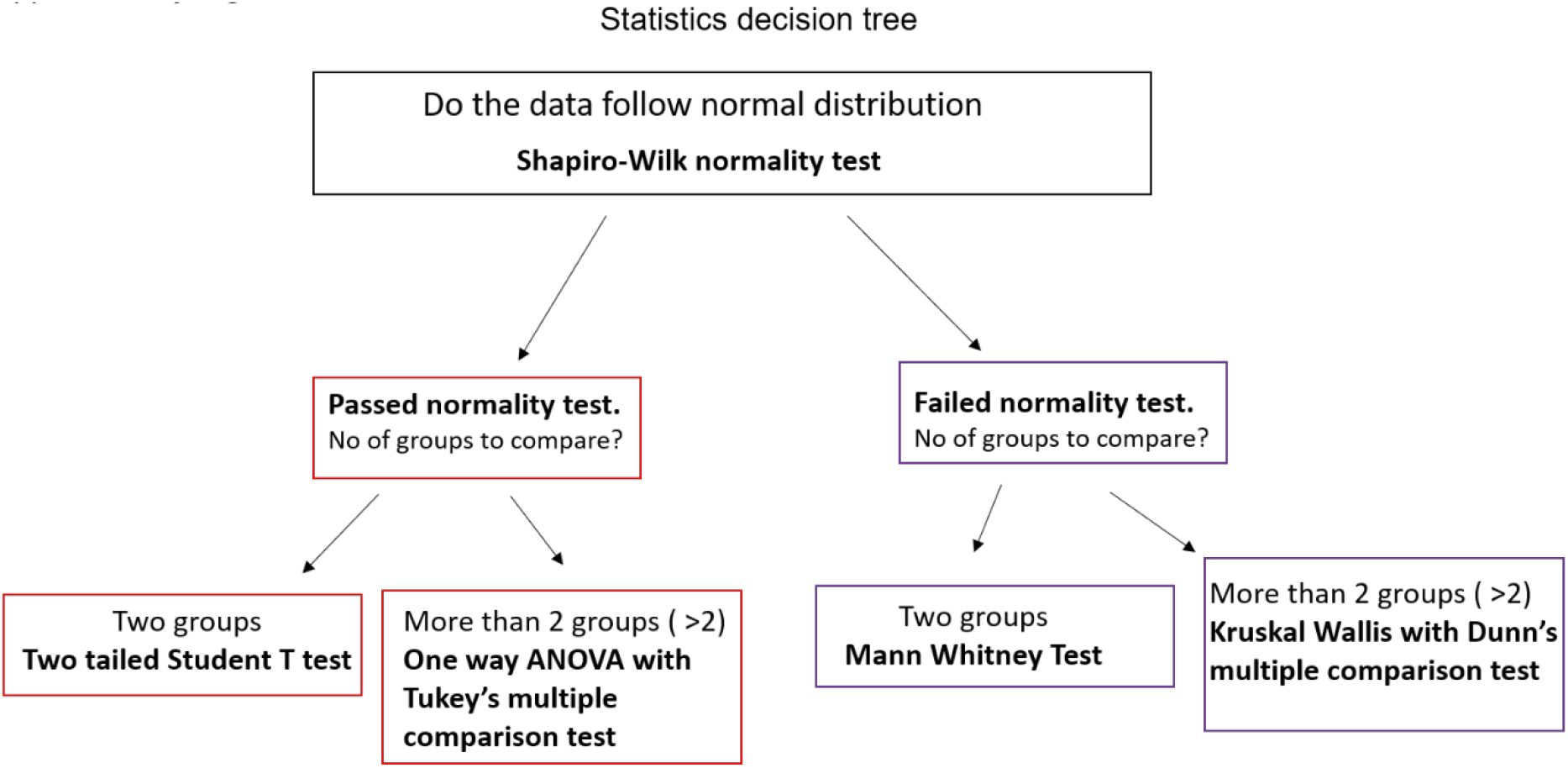
Statistical decision tree.

**Table S1:** Clinical and histopathological details of samples used for validation.

| ID | Sex | Age at presentation | Laterality | Clinical Risk | IIRC Group | AJCC staging |
| --- | --- | --- | --- | --- | --- | --- |
| P10 | F | 23 months | Bilateral | Advanced | Group E | cT3b |
| P11 | F | 24month | Unilateral | Advanced | Group E | cT3b |
| P12 | M | 36 months | Bilateral | Advanced | Group E | cT3b |
| P13 | F | 33 months | Unilateral | Advanced | Group E | cT3a |
| P14 | M | 7 months | Bilateral | Non-advanced | Group D | cT2a |
| P15 | M | 30 months | Bilateral | Non-advanced | Group D | cT2b |
| P16 | F | 14months | Unilateral | Non-advanced | GroupD | cT2b |
| P17 | M | 11 months | Unilateral | Non-advanced | Group D | cT2b |
| C1 | M | 2 months |  | Multiple organ dysfunction ( No ocular complications) |  |  |
| C2 | F | 12 months |  | No ocular complications |  |  |
| C3 | M | 3 months |  | No ocular complications |  |  |
| C4 | M | 6 months |  | No ocular complications |  |  |

